# Reliability of Whole-Exome Sequencing for Assessing Intratumor Genetic Heterogeneity

**DOI:** 10.1101/253195

**Authors:** Weiwei Shi, Charlotte K. Y. Ng, Raymond S. Lim, Tingting Jiang, Sushant Kumar, Xiaotong Li, Vikram B. Wali, Salvatore Piscuoglio, Mark B. Gerstein, Anees B. Chagpar, Britta Weigelt, Lajos Pusztai, Jorge S. Reis-Filho, Christos Hatzis

## Abstract

Multi-region sequencing is used to detect intratumor genetic heterogeneity (ITGH) in tumors. To assess whether genuine ITGH can be distinguished from sequencing artifacts, we whole-exome sequenced (WES) three anatomically distinct regions of the same tumor with technical replicates to estimate technical noise. Somatic variants were detected with three different WES pipelines and subsequently validated by high-depth amplicon sequencing. The cancer-only pipeline was unreliable, with about 69% of the identified somatic variants being false positive. Even with matched normal DNA where 82% of the somatic variants were detected reliably, only 36%-78% were found consistently in technical replicate pairs. Overall 34%-80% of the discordant somatic variants, which could be interpreted as ITGH, were found to constitute technical noise. Excluding mutations affecting low mappability regions or occurring in certain mutational contexts was found to reduce artifacts, yet detection of subclonal mutations by WES in the absence of orthogonal validation remains unreliable.

## INTRODUCTION

Intratumor genetic heterogeneity (ITGH), typically defined as the coexistence of genetically distinct but clonally related cancer cells within the same patient (Yap et al., 2012), can manifest itself spatially within the same lesion or as genetic differences between different metastatic sites and the primary tumor from the same patient (Ding et al., 2010; Gerlinger et al., 2012; Marusyk et al., 2012; Newburger et al., 2013; Yates et al., 2015). The broad availability of massively parallel sequencing has accelerated research into ITGH and numerous studies have applied whole-exome or targeted exome sequencing to multiple biopsies from the same cancer, to different metastatic lesions from the same patient, and more recently to multiple single cells from the same cancer (Gerlinger et al., 2012; Hou et al., 2012; Martelotto et al., 2017; Navin et al., 2011; Newburger et al., 2013; Nik-Zainal et al., 2012; Wang et al., 2014; Xu et al., 2012). ITGH represents a snapshot of the tumor’s evolutionary path and is a clinically important phenomenon with implications in prognosis and treatment response (Fisher et al., 2013; Hiley et al., 2014; Jiang et al., 2014; Marusyk et al., 2012; Morris et al., 2016; Turner and Reis-Filho, 2012).

The assessment of ITGH, by definition, involves the detection of subclonal, low frequency variants that are not uniformly present in all cancer cells, and is made possible by the availability of bioinformatics tools to detect low frequency somatic mutations with high sensitivity (Cibulskis et al., 2013; Koboldt et al., 2012; Saunders et al., 2012; Wilm et al., 2012). However, the presence of technical noise in sequencing data is well known (Li, 2014; Nakamura et al., 2011) and it is unclear whether genuine ITGH can be reliably distinguished from artifacts generated during library preparation, sequencing, and data processing (Qi et al., 2015; Smith et al., 2014). Understanding the signal to noise characteristics in these experiments is critical for the interpretation of ITGH.

Given the implications in prognosis and treatment response, ITGH is an important consideration in the clinical setting. Because the cost of WES and complex informed consent requirements, the inclusion of matching normal samples still represents a limitation in the sequencing of tumor samples in some large clinical trials (Shi et al., 2017). Whether subclonal mutations can be robustly identified in the absence of matching normal samples and whether pooled normal samples from unrelated individuals would serve as a reasonable control should be explored as alternative options for the assessment of ITGH in the clinical setting.

In this study, we aimed to assess the reliability of somatic variant detection from whole exome sequencing (WES) in the context of ITGH (**Fig. 1a, Fig. S1**). To address this question, we performed WES on DNA from biopsies obtained from three anatomically distinct regions of six primary breast cancers (6 × 3 biopsies) and from matched peripheral blood leukocytes. Additionally, to determine the background noise levels as a comparator for the assessment of ITGH, aliquots of the same DNA samples from six distinct biopsies were sequenced twice to generate technical replicates. We examined the reliability of somatic variant detection using three different analysis approaches, namely, using cancer WES data only, cancer data and WES from pooled unrelated (i.e. non-matched) normal samples, and WES data from cancer and matching normal tissue (i.e. blood). To generate the ‘gold standard’ benchmark dataset, we re-sequenced all somatic variants identified by at least one of the three somatic variant detection pipelines in at least one sample using high-depth targeted amplicon sequencing (i.e. AmpliSeq). Given the higher depth obtained by amplicon sequencing, we assessed true ITGH and estimated the frequency of false positives by the different detection pipelines. Finally, we evaluated the sequence patterns and context in which the artifactual mutations occurred to improve the specificity of WES for characterizing genuine ITGH.

**Fig. 1.**
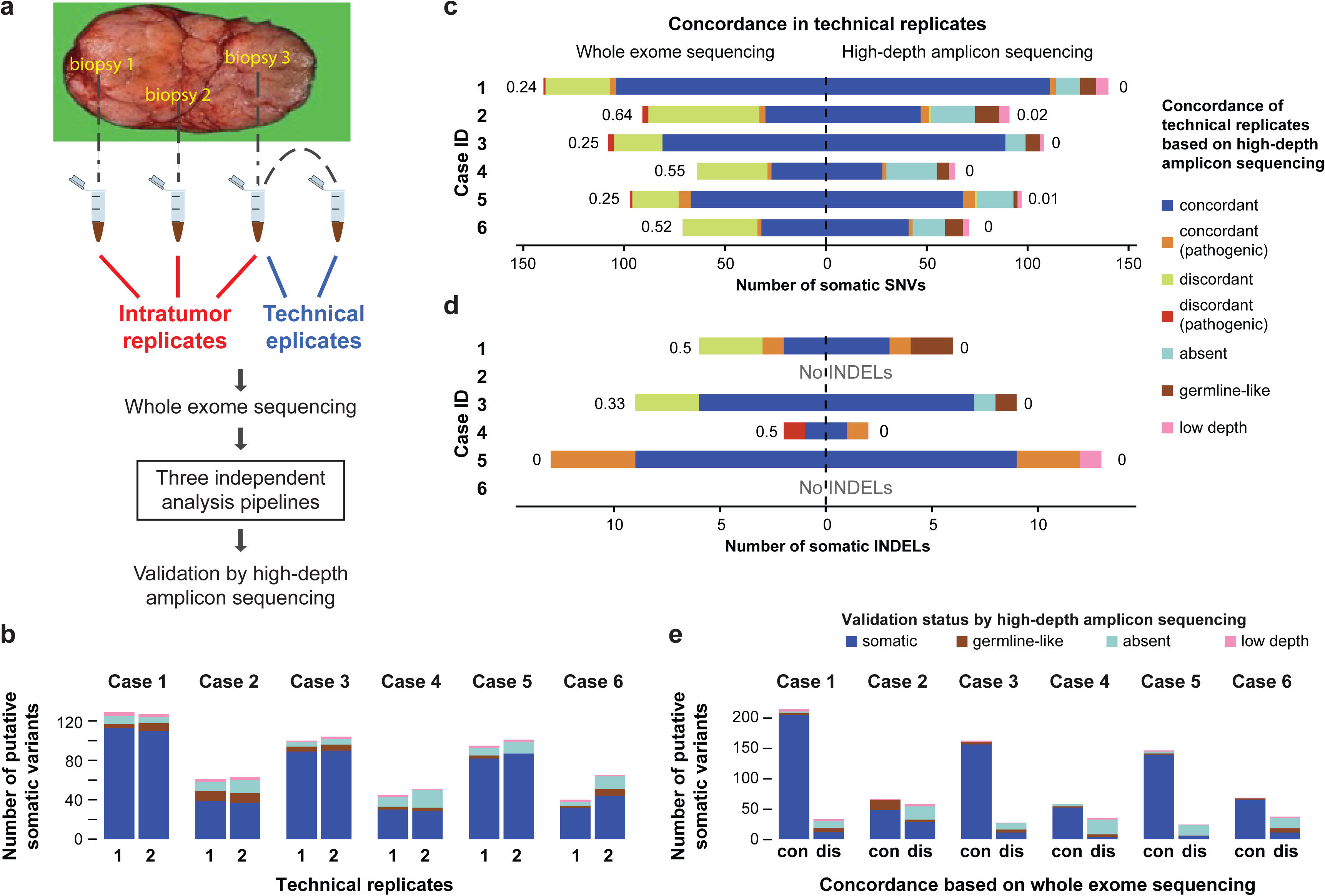
Reliability of somatic SNVs and INDELs detected by whole exome sequencing and high-depth amplicon sequencing validation of variants. **(a)** Biopsies were obtained from 3 anatomically distinct regions of each tumor to assess spatial genomic heterogeneity. One of the three DNA samples was split in two to provide a pair of technical replicates. Somatic variants were detected by three different WES analysis pipelines and subsequently validated by high-depth amplicon sequencing. (**b**) Number of somatic mutations identified by WES in each technical replicate, subclassified according to their validation status by high-depth amplicon sequencing. **(c-d)** Concordance of somatic SNVs (**c**) and INDELs (**d**) defined in each pair of technical replicates using the matched-normal WES analysis pipeline (left) or high-depth amplicon sequencing (right). Discordance between the replicates quantified as the Jaccard distance shown next to each bar and the pathogenicity of variants were assessed as described in the **Supplemental Experimental Procedures**. Putative WES variants re-sequenced with high-depth amplicon sequencing were further classified as absent (VAF<1%), germline (tumor VAF/germline VAF < 5), or low depth (<50x). **(e)** Validation status of variants (SNVs and INDELs) detected by WES in technical replicate pairs, categorized as concordant (“con”) or discordant (“dis”) by WES, and the distribution of their Ampliseq validation status is shown within each bar.

## RESULTS

### Limited reliability of somatic mutations defined by WES

We performed WES on DNA extracted from three distinct regions of the primary tumor and the matching normal blood cells from 6 breast cancer patients, including 4 with estrogen receptor positive and 2 with triple-negative cancers. The biopsies were obtained from 3 anatomically distinct regions of each tumor at least 1 cm apart (i.e. intratumor replicates, **Fig. 1a**) in the context of a prospective Institutional Review Board (IRB) approved study to assess intratumor molecular heterogeneity (von Wahlde et al., 2017). All tumor samples had at least 50% tumor cellularity based on pathologic assessment. For the technical replicates, a second library was generated from one of the tumor DNA samples randomly selected from each cancer and sequenced by WES using the same protocol at the same facility on a different day. The mean target depth was 160× (range 70× to 220×), consistent with recommendations for WES(Clark et al., 2011; Sims et al., 2014). Following WES, somatic single nucleotide variants (SNVs) and small insertion-deletions (INDELs) were identified by three different somatic variant calling pipelines that used either (i) the tumor DNA alone (i.e. tumor-only) (ii) tumor DNA and pooled unrelated normal DNA (i.e. cohort-normal) or (iii) tumor DNA and patient matched normal DNA (i.e. matched-normal, **Fig. S1** and **Supplemental Experimental Procedures**). Targeted amplicon sequencing using an orthogonal library generation and an independent sequencing method (AmpliSeq) was performed for all putative somatic variants identified by at least one of the WES variant calling pipelines on all tumor and matching normal DNA to a median depth of 605x to define the ‘gold standard’ mutation status for each identified somatic variant (**Supplemental Experimental Procedures**).

To quantify the technical reliability of somatic mutation detection by the matched-normal WES pipeline, the approach that is most frequently used in the research setting to assess ITGH, we compared the somatic mutations identified in the six pairs of technical replicates. In this experiment, tumor samples and normal samples were sequenced to a median coverage of 184× (range 92–211) and 90× (range 80–138), respectively. We identified a median of 74 (range 40–125) and 3 (range 0–13) somatic SNVs and INDELs, respectively, in each of the twelve DNA samples. Considering the high-depth amplicon sequencing results as the ‘gold standard’, we categorized candidate mutations detected in WES as true somatic, absent, germline-like (incorporating genuine germline variants and artifactual variant alleles caused by alignment biases and/or the sequencing technology)(Kim and Speed, 2013) or low-depth (i.e. technical failure with amplicon sequencing, **Table S2** and **Supplemental Experimental Procedures**). Excluding the low-depth (technical failure) variants, a median 82% (range 56%-90% per sample) of the somatic SNVs were confirmed as somatic, a median 5% (range 0%-16%) as germline-like and the remaining were absent by Ampliseq (**Fig. 1b)**.

Given that both technical replicates used the same input DNA, we anticipated detecting nearly identical somatic mutations in each pair of replicates. However, only a subset of the somatic SNVs and INDELs were consistently identified in technical replicate pairs, with median Jaccard distances (ranging from 0, perfect agreement, to 1, absence of overlapping variants; see **Supplemental Experimental Procedures**) of 0.39 (range 0.24-0.64) for SNVs (**Fig. 1c**) and 0.17 (range 0-0.50) for INDELs (**Fig. 1d**). Interestingly, the technical replicate pairs with the highest Jaccard distances were those with the lowest tumor cell content as inferred by FACETS (Shen and Seshan, 2016) (**Table S1**). There was also a small number of potentially pathogenic variants among the discordant variants in technical replicate pairs (range 0-3, red bars in **Fig. 1c**) that could have been misinterpreted as ITGH. We obtained fewer somatic mutations but similarly modest reproducibility with the cohort-normal WES pipeline, and far fewer somatic mutations but improved reproducibility using the tumor-only pipeline (see explanation below; **Fig. S2**). Comparing only the mutations that were confirmed to be somatic by Ampliseq between the technical replicate pairs, we observed almost perfect agreement for SNVs (**Fig. 1c**) and for INDELs (**Fig. 1d**) (maximum Jaccard distance of 0.02 and 0, respectively).

When we examined the somatic mutations found to be concordant or discordant between pairs of technical replicates based on WES, a median of 95% (range 73%-97%) of the concordant variants were confirmed to be genuinely somatic in at least one of the two technical replicates, compared to a median 33% (range 14%-48%) of the discordant variants (**Fig. 1e**). Of the discordant WES variants, a median of 44% (range 36%-71%) were found to be absent by Ampliseq (i.e. false positive in one of the two technical replicates), and a median of 7% (range 3%-22%) were germline(-like) variants (i.e. missed by WES in the matching normal). The validation status of 3% (range 1% to 8%) of the mutations could not be ascertained due to technical failure in the validation experiments due to low Ampliseq coverage.

Taken together, these results suggest that WES performed at typical sequencing depth may be inadequate for detecting ITGH, particularly when the tumor cell content is less than 50%, as only 62% (range 36% to 76%) of the somatic mutations were detected consistently in the technical replicate pairs by WES with the remaining mutations falsely appearing as discordant.

### WES overestimates true intratumor genetic heterogeneity

Next, we quantified ITGH by comparing somatic variant calls between the geographically distinct biopsies from the same cancer. In this analysis, we also examined how the three different WES analytic approaches differed in the quantification of ITGH (**Fig. S1** and **Supplemental Experimental Procedures**). Tumor cellularity of all samples was inferred from WES using FACETS (Shen and Seshan, 2016) (**Table S1**). We used mixed-effects linear modeling to estimate an average cellularity of 54.7% with intratumor and technical standard deviation of 7.0% and 2.2%, respectively (**Supplemental Experimental Procedures)**. The matched-normal pipeline detected a median of 150 unique somatic mutations (range 68-186) in all 3 samples combined per tumor. Compared to the matched-normal pipeline, the cohort-normal pipeline detected a median of 101 mutations (range 48-131, Wilcoxon test n.s.) and the tumor-only pipeline identified a median of 62 mutations (range 36-97, Wilcoxon test p=0.01) (**Fig. 2a–b**).

**Fig. 2.**
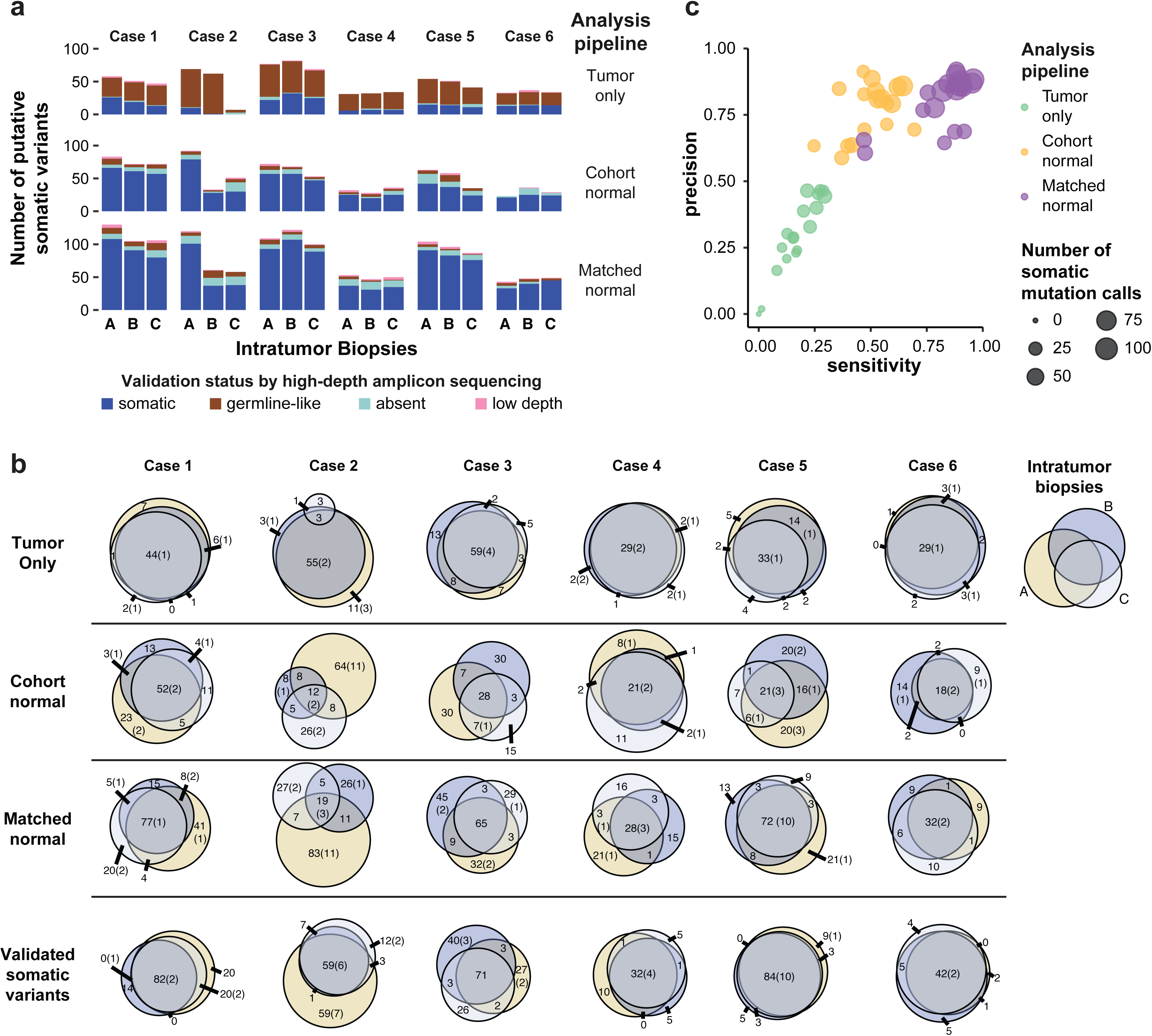
Intratumor genetic heterogeneity as assessed by different WES analysis pipelines. **(a)** Total number of somatic variants (SNVs and INDELs) identified in intratumor biopsies by the three WES analysis pipelines. One of the two technical replicates was randomly selected for inclusion in this analysis. Validation status by high-depth amplicon sequencing (i.e. somatic, germline, absent [VAF < 1%], low depth [< 50x coverage]) is shown according to the color key. **(b)** Venn diagrams showing the overlap of putative somatic variants detected in intratumor biopsies from each tumor by the three WES analysis pipelines. The last row includes only “true” somatic variants validated by high-depth amplicon sequencing. The size of the circles is proportional to the number of somatic variants in a biopsy, with the numbers representing the total variants and those in parentheses indicating the number of pathogenic variants. **(c)** Performance characteristics of the three WES analysis pipelines to identify true somatic variants. Putative somatic variants were considered as “true” if confirmed by high-depth amplicon sequencing. Precision was calculated as TP/(FP+TP) and sensitivity as TP/STP, where TP and FP are the number of true positive (TP) and false positive (FP) variants and STP is the total number of true somatic calls made by all three pipelines. Each circle represents one sample as analyzed by each pipeline, and the size of the circles is proportional to the number of putative somatic variants per biopsy identified by each analysis pipeline.

To assess the reliability of the different WES pipelines in detecting ITGH, we compared the WES candidate mutations to the ‘gold-standard’ Ampliseq validated somatic variants from the same sample. A median of 62% (range 50%-98%) of the candidate somatic variants detected by the tumor-only pipeline were germline variants, and only 28% (range 2%-45%) validated as true somatic mutations. By contrast, a median of 79% (range 59%-91%) and 84% (range 61%-92%) of the variants defined by the cohort-normal and the matched-normal pipelines, respectively, were true somatic mutations (**Fig. 2a**, **Fig. S3a-b**). The tumor-only pipeline had the lowest sensitivity (median 18%, range 0%-29%) and precision (median 28%, range 0%-46%) for identifying true somatic variants. The matched-normal pipeline had the highest sensitivity (median 87%, range 47%-94%) and precision (median 86%, range 62%-94%, **Fig. 2c**), whereas the cohort-normal pipeline had similar precision (median 83%, range 61%-91%) but significantly lower sensitivity (median 48%, range 26%-64%; Wilcoxon p < 0.001).

Next, we estimated the apparent ITGH based on mutations detected by the three WES calling pipelines using the Jaccard distance as the metric of ITGH. Mutations identified as somatic in one or two of the three biopsies and as germline-like or absent in the remaining biopsies contributed to ITGH. The tumor-only pipeline had a median Jaccard distance of 0.34 (range 0.19-0.96) compared to the cohort-normal pipeline of 0.70 (range 0.53-0.91) and matched-normal pipeline of 0.60 (range 0.44-0.89, **Fig. 2b**). The apparently lower ITGH defined by the tumor-only pipeline (paired Wilcoxon test p=0.03 for tumor-only vs cohort-normal; ns vs matched-normal) was due to the high number of germline variants misidentified as somatic mutations by the tumor-only pipeline (**Fig. 2a**). When ITGH was estimated based on the Ampliseq validated somatic mutations only, the median Jaccard distance was 0.40 (range 0.19-0.61, **Fig. 2b**), which in the context of this study was considered a true estimation of ITGH. Compared to the apparent ITGH defined by the candidate mutations in the cohort-normal and matched-normal WES pipelines, the true ITGH was significantly smaller (paired Wilcoxon test p=0.015 vs cohort-normal and p=0.015 vs matched-normal). We noted that 5.8% of the heterogeneous variants, also called branch mutations that were not present in all biopsies of a given case, were predicted to be pathogenic, but there was no statistically significant enrichment in pathogenic mutations compared to non-pathogenic variants among the heterogeneous somatic variants (**Fig. 2b**). Heterogeneous variants were detected in a small number of cancer genes (21 of 306 or 6.9%) and 5 of these variants (1.6%) were predicted pathogenic.

Taken together, these results suggest that the tumor-only WES pipeline misidentifies a substantial proportion of germline variants as somatic mutations. Even when using the matched-normal DNA for mutation detection, the extent of ITGH defined based solely on WES performed at typical sequencing depth is overestimated, potentially affecting actionable cancer genes. For example, a deleterious stop-gain branch mutation in *CDC27* (p.Cys71*) was identified as heterogeneous (in one of the 3 biopsies) by the matched-normal WES pipeline, but was not validated by Ampliseq. False positive heterogeneous variants were mostly not actionable, however.

### Characteristics of artifactual WES somatic mutations

To identify the characteristics of the putative mutations identified by WES that were subsequently confirmed not to be truly somatic variants, we examined the alternative coverage (i.e. the number of reads supporting the alternate allele), the variant allele frequency (VAF) and the total depth of coverage of the candidate somatic mutations identified by the three WES pipelines from the intratumor biopsies. The tumor-only pipeline reported a median of 2 variants (range 0-5) with VAF<10% due to reduced sensitivity of the single-sample mutation detection algorithm at low VAF and the strict filtering imposed to remove potential germline variants (**Fig. 3a–b**). Despite aggressive filtering, most of the putative somatic variants from the tumor-only WES pipeline were found to constitute germline variants with VAF~50%, indicating the presence of a large number of private mutations not having been catalogued in publicly available databases (**Figs. 2a and 3a–b**). The cohort-normal WES pipeline correctly identified subclonal somatic variants with low VAF, including many that were heterogeneous between the biopsies, but missed somatic variants with VAF>45% due to filtering imposed to remove likely germline variants that may not be present in the pooled normal DNA used as the reference (**Figs. 2c and 3a–b**). The validated somatic mutations defined by the matched-normal pipeline covered the widest range of VAFs. The putative somatic mutations identified by the matched normal pipeline that were confirmed to be absent by Ampliseq were mostly in the low VAF range (median 7.3%, range 0.6%-44%, **Figs. 2a and 3a–b**).

**Fig. 3.**
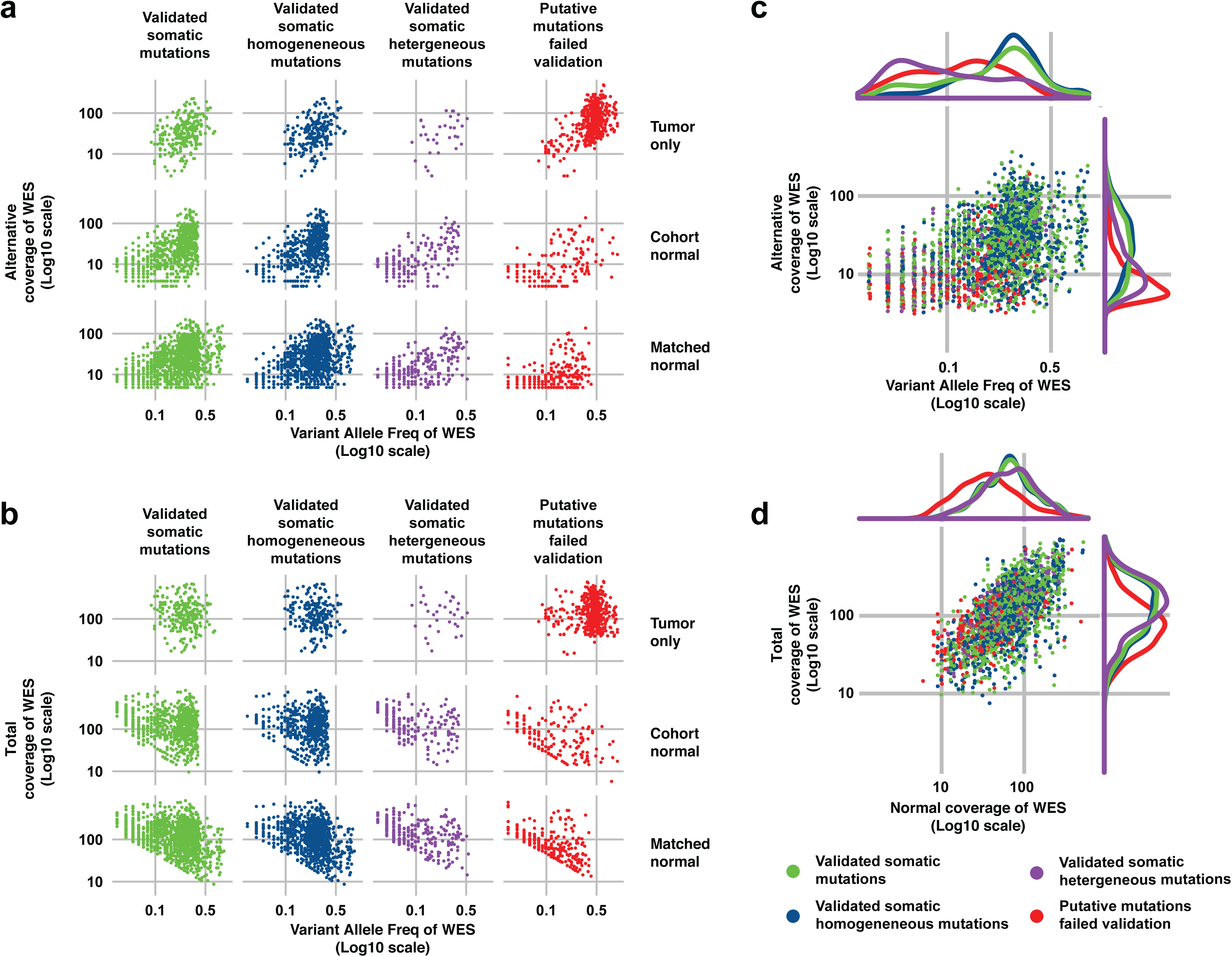
Coverage characteristics of true somatic variants and false positive mutations in the WES data. **(a)** Alternative allele coverage (i.e. number of read supporting the alternative allele) and **(b)** total coverage are plotted against variant allele fraction (VAF, all log10 scale) for the three WES analysis pipelines (rows). The different subsets of WES putative somatic variants according to validation status by high-depth amplicon sequencing are shown as columns: validated somatic mutations, validated homogeneous somatic mutations (i.e. present in all three biopsies of the same tumor), validated heterogeneous somatic mutations (i.e. present in one or two biopsies from the same tumor), and putative somatic mutations identified by the WES pipelines but failed validation by high-depth amplicon sequencing (i.e. putative somatic mutations that were validated to be germline, absent [VAF < 1%] or low coverage [< 50x]). For the matched-normal WES pipeline, **(c)** WES alternative allele coverage (all log10 scale) is plotted against VAF and (**d**) total coverage in the tumor (bottom) is plotted against the coverage in the matched normal sample of somatic mutations identified in all the specimens. The validation status categories are the same as in panels (a) and (b). Density kernel plots of the marginal distributions are included above and to the right of the scatter plots for each of the four categories of mutations.

Given that the majority of the ITGH studies carried out to date (Ding et al., 2010; Gerlinger et al., 2012; Nik-Zainal et al., 2012) used matched tumor-normal samples and analysis pipelines similar to our matched-normal pipeline and that mainly low VAF mutations contribute to ITGH (**Fig. 3a–b**), we compared the true and false positive somatic mutations (i.e. the validated and the unvalidated putative somatic variants) and the validated homogeneous (i.e. present in all biopsies from a case) and heterogeneous (i.e. absent in at least one biopsy from a case) somatic mutations derived from the distinct biopsies using the matched-normal WES pipeline. We found that the total depth for the true positive (median 124, range 9-1028, green in **Fig 3c**) was significantly higher than for the false positive mutations (median 78, range 12-926, p<0.001, Wilcoxon test, red in **Fig 3c**). Furthermore, the false positives, compared to true mutations, had significantly lower alternative coverage (median 7, range 1-126 vs median 22, range 4-231, p<0.001, Wilcoxon test) and VAF (median 11%, range 0.6%-64% vs median 23%, range 3%-93%, p<0.001, Wilcoxon test, **Fig 3c**). There were also significant differences in the VAF distribution of the validated homogeneous and heterogeneous mutations (median 25%, range 1%-93% vs median 6%, range 1%-57%, p<0.001, Wilcoxon test) and between the validated homogeneous mutations and the false positives (median 25%, range 1%-93% vs median 12%, range 2%-64%, p<0.001, Wilcoxon test, **Fig 3c**). Crucially, the validated mutations implicated in true ITGH had significantly lower VAF than the false positives (p<0.001, Wilcoxon test, **Fig. 3c)**, **Fig. S4**), which indicates that true ITGH mutations may display similarly low or lower VAFs as the false positive and false negative mutations. These results suggest that filtering somatic variants with low VAF or low alternative coverage may improve the precision of the WES pipeline but would also eliminate many true somatic variants that contribute to ITGH. Importantly, false positive mutations have significantly lower total depth in the matched normal DNA (median 35, range 6-492) compared to true positive mutations (median 68, range 9-514, p<0.001, Wilcoxon test) and validated heterogeneous mutations (median 72, range 12-343, p<0.001, Wilcoxon test, **Fig. 3d)**. Many putative somatic mutations (47%) with total depth in the normal DNA of 10 or less were confirmed as germline-like by Ampliseq, emphasizing the importance of having adequate sequencing depth in the normal sample as well.

### Separating true ITGH from WES artifacts

Because subclonal mutations are expected to be the predominant contributors to ITGH, we inferred the clonality of all mutations identified by WES using ABSOLUTE (Carter et al., 2012). Subclonal mutations were significantly overrepresented among the validated heterogeneous variants compared to the homogeneous somatic variants (83.2% vs 28.9%, p<0.001, Fisher exact test, **Fig. 4a**), but were similarly overrepresented among the artifactual somatic variants (91.5% vs 83.2%, n.s., **Fig. 4a**). Importantly, subclonal mutations were also significantly enriched among discordant variants in the WES technical replicates in compared to the concordant variants (72.0% vs 24.9%, p<0.001, Fisher exact test, **Fig. 4a**). These results suggest that, compared to clonal variants, subclonal variants detected by WES are more likely to be erroneously attributed to ITGH.

**Fig. 4.**
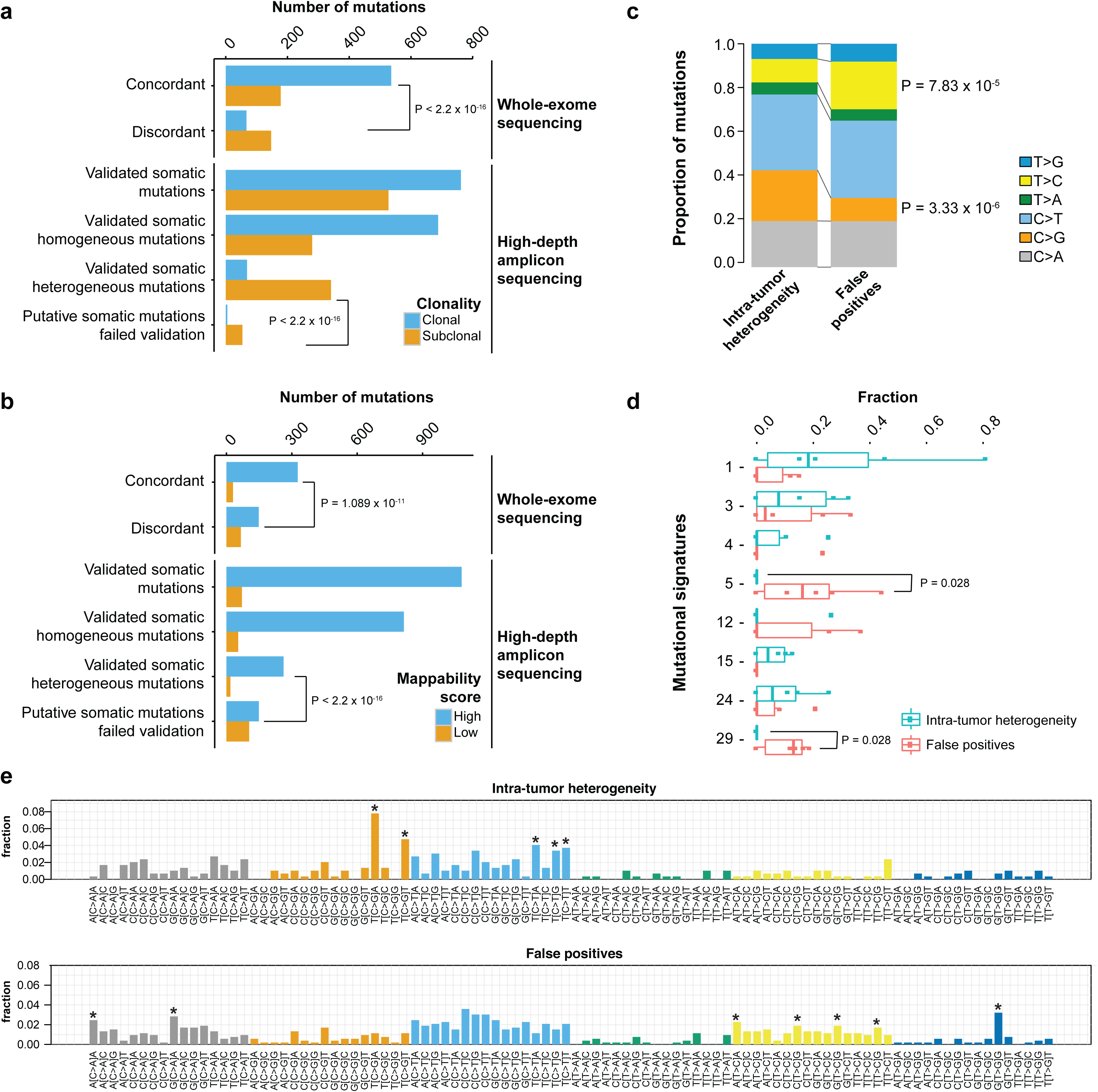
Mappability and sequence context of true and artifactual somatic variants. **(a)** Clonality as defined by ABSOLUTE and (**b**) mappability of somatic variants identified by the matched-normal WES pipeline. In each panel, the first two sets of bars enumerate the putative somatic variants identified as concordant or discordant in the technical replicates, whereas the bottom four sets of bars enumerate the somatic variants identified in intratumor biopsies and subsequently validated by high-depth amplicon sequencing. High mappability regions are regions with mappability score of 1 (see **Supplemental Experimental Procedures**). **(c)** Comparison of the mutational spectra of validated somatic heterogenous mutations and artifactual somatic mutations that failed validation in all samples. The reference base listed (C or T) includes the corresponding reverse complement (G or A). **(d)** Distribution of signature weights obtained from the decomposition of mutational signatures from each tumor sample. **(e)** Detailed mutational spectra of the trinucleotide context of the pool of mutations detected in all tumor samples. Trinucleotide contexts with significant enrichment in the validated somatic heterogeneous mutations or in the artifactual somatic mutations are shown with an asterisk above the corresponding bars. Asterisks indicate p < 0.005. Statistical comparisons in (**a**), (**b**), (**c**) and (**e**) are based on Fisher’s exact tests. Statistical comparisons in (**d**) are based on Wilcoxon tests. All statistical tests are two-sided. p<0.05 was considered statistically significant.

Examination of the sequence context of mutations revealed that 41.1% of the artifactual somatic mutations occurred in genomic regions of low mappability (Derrien et al., 2012) compared to only 6.4% for the validated somatic heterogeneous mutations (p<0.001, Fisher exact test, **Fig. 4b**). Furthermore, 30.8% of the discordant variants in the WES technical replicates occurred in low mappability regions compared to 8.4% for the concordant variants (p<0.001, Fisher exact test, **Fig 4b**). These results suggest that ambiguous mapping of DNA fragments directly contributes to artifactual somatic variants. Compared to the validated heterogeneous mutations, artifactual somatic mutations appeared to be significantly enriched in T>C transitions (p<0.001, Fisher exact test, **Fig. 4c**), particularly in the ApTpA and NpTpG trinucleotide contexts and also in T>G transversions in the GpTpG context (**Fig. 4e**). By contrast, artifactual somatic mutations were significantly depleted in C>G transversions (p<0.001, Fisher exact test, **Fig. 4c**). Interestingly, validated somatic heterogeneous mutations were enriched for C>G and C>T in the TpCpA and TpCpT contexts (**Fig. 4e**), the characteristic substitution patterns induced by the upregulation of APOBEC cytidine deaminases (Nik-Zainal et al., 2016). Although a substantial proportion of the artifactual mutations detected were C>T transitions, which have been associated with both the aging process and with lab-induced cytosine deamination during DNA library preparation (Alexandrov et al., 2013; Chen et al., 2014), their proportion was similar between the artifactual and validated somatic heterogeneous variants (34.6% vs 33.9%, n.s., **Fig. 4c**), with mutations occurring in the TpCpN context being significantly underrepresented in the artifactual variants (**Fig. 4e**). Mutational signature analysis using previously defined mutational signatures(Alexandrov et al., 2013) identified Signatures 5 and 29 to be overrepresented in the artifactual somatic mutations (p=0.028, Wilcoxon test, **Fig. 4d**), with a median of 16.1% (range 0%-44.4%) of the artifactual mutations classified as Signature 5, mainly driven by the T>C transitions reported above. Signature 29 is driven by C>A mutations, predominantly occurring in the ApCpA and GpCpA contexts that were significantly overrepresented among the artifactual variants (**Fig. 4e**).

Finally, we considered whether applying filtering strategies to exclude mutations that occur in low mappability regions or within sequence contexts which are enriched in artifactual mutations (C>A mutations in ApCpA and GpCpA contexts, T>C mutations in ApTpA and [C/T/G]pTpG contexts, and T>G mutations in GpTpG context, **Fig 4e**) can improve the reliability of WES. Excluding putative somatic mutations in low mappability regions improved the precision for mutations detected in the technical (0.862 vs 0.792 without filtering) and biological replicates (0.876 vs 0.817 without filtering), while reducing moderately the sensitivity (0.858 vs 0.913 for technical; 0.773 vs 0.824 for biological). Filtering by mutational context reduced the sensitivity without appreciably improving precision, as did filtering by the combination of filters (**Fig. S5).**

## DISCUSSION

WES is an appealing and increasingly affordable technology to study the extent of ITGH in an unbiased manner (i.e. without a priori selecting genes of interest for sequencing). The reliability of WES to detect low frequency mutations, which often account for the majority of ITGH within a cancer, depends on the experimental design, the sequencing depth and the bioinformatics approaches used to define the somatic variants. To examine the influence of these factors on measuring ITGH, here we generated a dataset incorporating both technical and biological replicates sequenced to depths commonly found in clinical or translational research studies, and validated every putative somatic variant detected with orthogonal high-depth sequencing methods. All datasets generated in this study have been made publicly available to provide a resource for the community to refine analytical tools for ITGH detection from WES.

We performed six pairs of technical replicates that involved independent library preparation and sequencing of aliquots from the same DNA extractions, and experiments were performed on different days. The technical replicates revealed an unexpectedly high degree of discordance in the putative somatic variants identified, even using the current best practice matched-normal variant calling analysis approach. Subsequent validation with high-depth amplicon sequencing (605x median coverage) of all variants identified by WES demonstrated that the majority of the false positive somatic variants either i) displayed low variant allele fraction and were often detected as subclonal in one experiment but not in the other, ii) were in fact germline-like variants that appeared as heterogeneous somatic mutations (Kim and Speed, 2013), or iii) map to genomic regions of low mappability. The enrichment of subclonal mutations among the discordant mutations in the WES technical replicates is expected, given the well known difficulty in identifying somatic mutations at low VAF. Indeed, comparison of the putative mutations that did not validate to the ‘true’ somatic mutations that were validated by high-depth amplicon sequencing demonstrated that mutations with low VAF and/or low alternative coverage were more difficult to be reliably identified by WES. On the other hand, our results revealed a not insignificant proportion of germline-like false positive mutation calls. While some of these germline-like variants are genuinely germline alleles not detected in the matched normal samples, a substantial proportion of these are likely attributed to alignment and sequencing biases (Kim and Speed, 2013) that manifested as false positive variants at low VAF/alternative coverage. Importantly, our results highlighted the often overlooked importance of the coverage of the matched normal sample in the accurate identification of somatic mutations, given that mutations that failed validation, on average, were associated with lower coverage in the normal sample. In terms of mappability of genomic regions, we found that false positive mutation calls were enriched in genomic regions of poor mappability. In fact, we demonstrated that this may represent a reasonable filter if specificity is of paramount importance and some trade-off in sensitivity can be tolerated. Although our study provides direct evidence in support of ITGH, the mutations implicated in ITGH show substantial overlap with the alterations stemming from intrinsic technical noise associated with WES in terms of VAF, alternate allele depth, total depth in tumor and normal, as well as mappability.

Our analysis of the mutational signatures between the validated mutations implicated in ITGH versus the false positive mutations revealed striking differences. The heterogeneous mutations were enriched in a pattern typically associated with increased APOBEC activity and this pattern has been previously shown to contribute to ITGH in breast and lung cancers (de Bruin et al., 2014; Ng et al., 2017). On the other hand, the false positive mutations were enriched for, in particular, C>A and T>G mutations at specific sequence contexts. A recent study of rare polymorphisms determined by high-depth whole genome sequencing in 300 individuals of diverse genetic origins identified four mutational signatures, two of which were consistent within populations and had a clear association with geographic distribution (Mathieson and Reich, 2017). The origin of the remaining two were uninterpretable, with one of these latter signatures dominated by T>G mutations in the GpTpG context, and the other signature highly correlated with COSMIC Signature 5, which has been found in all cancer types (Alexandrov et al., 2013) and has been suggested to display clock-like properties suggestive of an association with the aging process (Alexandrov et al., 2015). Our analysis of the sequence context of false positive variants identified both these features as being significantly enriched in artifactual putative mutations (**Fig. 4c–d**), strongly suggesting that caution should be exercised in the interpretation of the reported mutational signatures.

This study has several limitations. The sample size of 6 breast cancers and 18 biopsies may be too limited to allow the generalization of our results on ITGH to all breast cancer subtypes and to other cancers. Of note, the unique nested experimental design incorporating within-sample technical replication processed and sequenced in the same manner as the intra-tumor biopsies, which provided an estimate of background discordance against which the ITGH results could be interpreted. Additionally, the extensive orthogonal validation by high-depth amplicon sequencing on an independent sequencing platform with very different chemistry from the platform used for WES adds rigor to our study. The sequencing depth attained in this study is comparable to previous studies utilizing WES (with subsequent high-depth sequencing for validation) for the genomic characterization of ITGH (Yates et al., 2015); it is plausible, however, that WES at higher depth (i.e. >250x) would mitigate in part the false positives and false negatives, in particular in samples with tumor cell content <50%.

In summary, our study showed that WES at 184 mean depth of coverage in the tumor samples overestimates the extent of ITGH, and the technical noise associated with somatic mutation detection using WES alone can confound true ITGH. Our results also suggested that it is not possible to reduce the false positive rate through more aggressive minimum depth filtering without impacting the sensitivity of detecting true somatic mutations in the 1%-5% VAF range, but excluding mutations that occur in low mappability regions of the genome, or in certain mutational contexts could reduce artifactual somatic mutations and provide less biased estimates of ITGH. Nevertheless, orthogonal, high-depth validation experiments are highly desirable in the context of quantifying ITGH.

## ACKNOWLEDGEMENTS

Research reported in this publication was supported in part by the Breast Cancer Research Foundation (to LP, JSR-F and CH) and by a Cancer Center Support Grant of the National Institutes of Health/National Cancer Institute (grant No P30CA008748). SP is currently funded by Swiss National Foundation (Ambizione grant number PZ00P3_168165). The content is solely the responsibility of the authors and does not necessarily represent the official views of the National Institutes of Health. The funders had no role in study design, data collection and analysis, decision to publish, or preparation of the manuscript.

## AUTHOR CONTRIBUTIONS

Conceptualization, L.P., J.S.R.-F., B.W. and C.H.; Methodology, L.P., J.S.R.-F., W.S., C.K.Y.N., M.B.G., and C.H.; Software, W.S., C.K.Y.N., R.S.L., T.J., S.K., and X.L.; Validation, C.K.Y.N., R.S.L., B.W. and J.S.R.-F.; Formal Analysis, W.S., C.K.Y.N., R.S.L., T.J., S.P. and C.H.; Investigation, V.B.W., A.B.C., and B.W.; Resources, L.P., A.B.C., M.B.G., B.W., J.S.R.-F. and C.H.; Data Curation, V.B.W. and A.B.C.; Writing – Original Draft, W.S., C.K.Y.N., R.S.L., B.W., J.S.R.-F. and C.H.; Writing – Review and Editing, all authors; Visualization, W.S., C.K.Y.N., R.S.L., X.L. and C.H.; Supervision, L.P., J.S.R.-F., B.W., M.B.G. and C.H.; Funding Acquisition, L.P., C.H., J.S.R.-F.

## EXPERIMENTAL PROCEDURES

### Tumor sample collection

Breast cancer samples were collected from patients with newly diagnosed invasive breast cancer with tumor size > 2 cm at the Yale Cancer Center. Tumor tissues were obtained with 3 punch biopsies at least 1 cm apart from 3 different regions of the tumor after pathologic gross examination. Six of these tumors with high enough cellularity (>50%) and high DNA quality from all three biopsies and with matched blood DNA were selected for this study.

### Whole exome sequencing and analysis

DNA was extracted and library was prepared using standard protocols, and the exome was captured using the NimbleGen SeqCap EZ Human Exome Kit v2.0. Sequencing was performed on the HiSeq 2000 in paired-end 75-cycles mode at the Yale Center for Genome Analysis. We used three different analytical pipelines for detecting variants. A single-sample “tumor-only” pipeline, a “cohort-normal” pipeline suing an in-house normal reference obtained from 10 unrelated normal blood DNA samples, and a “matched normal” pipeline using the matched normal DNA from each patient as reference. Further details are provided in **Supplemental Experimental Procedures.**

### Validation of putative variants with high-depth amplicon sequencing

Variants identified by WES were subjected to validation with high-depth amplicon sequencing using custom Ampliseq panels on the same tumor and matched normal DNA samples. Amplicon sequencing was performed to a median depth of 600x.

### Mutational signature analysis and mappability

Mutational signature analyses comparing the validated variants that contribute to ITHG and the WES false positive calls were performed for individual tumor samples and for the pooled mutations over all samples using the R package deconstructSigs(Rosenthal et al., 2016). Mappability of SNVs was assessed using the CRG Alignability track(Derrien et al., 2012) in the UCSC genome browser. Additional details are in **Supplemental Experimental Procedures.**

### DATA AND SOFTWARE AVAILABILITY

All the raw data from WES and Ampliseq sequences generated in this project were deposited in the Sequence Read Archive (<u>https://www.ncbi.nlm.nih.gov/sra</u>) under accession ID SRP070662.

## SUPPLEMENTAL INFORMATION

Supplemental information includes Supplemental Experimental Procedures, five figures and five tables and can be found with this article online at

